# Genetically encoded nAChR upregulation is neuroprotective in female parkinsonian mice

**DOI:** 10.1101/2025.06.23.661175

**Authors:** Gauri Pandey, Roger C. Garcia, Debanjana Das, Donovan Pollock, Nethra Karthik, Sushmitha Nalluri, Tan Nguyen, Christopher Polo, Sara M. Zarate, Rahul Srinivasan

## Abstract

Parkinson’s disease is projected to rise to pandemic proportions by 2050, which has resulted in an urgent need for disease-modifying treatments. In this regard, we previously showed that in a mouse model of parkinsonism with unilateral 6-hydroxydopamine (6-OHDA) injection into the dorsolateral striatum (DLS), low doses of the neuronal nicotinic acetylcholine receptor (nAChR) partial agonist and smoking cessation drug, cytisine exerts sex-specific neuroprotection in substantia nigra pars compacta (SNc) dopaminergic (DA) neurons of only female mice by reducing apoptotic endoplasmic reticulum (ER) stress. Although these data suggest that neuroprotection might occur via cytisine-mediated upregulation of β2 subunit-containing (β2*) nAChRs in SNc DA neurons, there is no direct evidence to support this idea. Therefore, this study asks the critical question of whether upregulation of β2* nAChRs in SNc DA neurons alone is sufficient to reduce apoptotic ER stress and exert neuroprotection in a preclinical unilateral DLS mouse model of 6-OHDA-induced parkinsonism. To address this question, we generate and characterize a novel β2-upregulated transgenic mouse line. These transgenic mice possess mutations in the M3-M4 intracytoplasmic loop of β2 subunits that cause constitutive upregulation of β2* nAChRs without nicotinic ligands. Surprisingly, when compared to wild-type littermates, only female β2-upregulated transgenic mice demonstrate upregulation of β2* nAChRs in SNc DA neurons as assessed by significant increases in Sec24D-containing ER exit sites (Sec24D-ERES). Using the optogenetic calcium and dopamine sensors, GCaMP6f and GRABDA respectively, we found significant increases in dihydro-beta-erythroidine (DhβE)-sensitive β2* nAChR-mediated calcium influx in SNc DA neuron dendrites and DhβE-sensitive acetylcholine (ACh)-evoked dopamine release at SNc DA neuron terminals of the DLS in female transgenic mice. We then used four independent readouts to assess neuroprotection of SNc DA neurons following unilateral 6-OHDA injection into the DLS, *viz*., contralateral apomorphine-induced rotations, preservation of SNc DA neurons, inhibition of a major proapoptotic ER stress protein, C/EBP homologous protein (CHOP) and glial fibrillary acid protein (GFAP) expression in SNc astrocytes. In all four readouts, female β2-upregulated transgenic mice showed significant neuroprotection. From a clinical perspective, this study shows that upregulation without nicotinic ligand-mediated activation of β2* nAChRs in SNc DA neurons can be a translationally viable disease-modifying strategy for Parkinson’s disease. In addition, we envision that the novel transgenic β2-upregulated mice created in this study will provide a valuable tool for understanding the role of nAChR upregulation in major neurological disorders such as addiction, anxiety, depression and dementia.

## Introduction

Parkinson’s disease has emerged as the fastest growing neurological disorder with a projected worldwide prevalence of 25 million by 2050.^1^ Despite this alarming report, current treatments are merely symptomatic and do not address the proximate cause of movement disorder in Parkinson’s disease, which is a rapid loss of substantia nigra pars compacta (SNc) dopaminergic (DA) neurons.^2^ In this context, we know that the accelerated loss of SNc DA neurons during Parkinson’s disease involves multiple pathogenic factors including misfolded α-synuclein, hyperoxidative stress and mitochondrial dysfunction.^3^ These multiple cellular insults converge on an apoptotic endoplasmic reticulum (ER) stress response in SNc DA neurons, consequently leading to neurodegeneration.^4–7^ It therefore follows that mitigating apoptotic ER stress, which is a major convergent pathway causing the death of SNc DA neurons, can provide a powerful disease-modifying treatment strategy for Parkinson’s disease.

We previously showed that in the 6-hydroxydopamine (6-OHDA) preclinical mouse model of parkinsonism, systemic treatment with low doses of the smoking cessation drug cytisine mitigates apoptotic ER stress and exerts neuroprotection only in female parkinsonian mice.^8^ We also showed that cytisine-mediated neuroprotection in female parkinsonian mice requires 17-β-estradiol because the depletion of systemically circulating 17-β-estradiol via two distinct manipulations, *viz.* ovariectomy or letrozole treatment abolished the neuroprotective effect of cytisine.^9^ Furthermore, systemic administration of exogenous 17-β-estradiol rescued cytisine-mediated neuroprotection in 6-OHDA-exposed female mice with systemic 17-β-estradiol depletion.^9^ Taken together, these findings suggest that the smoking cessation drug cytisine acts in combination with systemically circulating 17-β-estradiol to exert neuroprotection against parkinsonism in female mice.^8,9^ Indeed, cytisine is being considered as a potential clinical treatment option for a number of neurological disorders.^10^

Importantly, our finding that low-dose cytisine exerts neuroprotection in female parkinsonian mice is predicated on the ability of low-dose cytisine to chaperone β2* containing neuronal nicotinic acetylcholine receptors (nAChRs) (*denotes the presence of other uncharacterized nAChR subunit subtypes in receptor pentamer) out of the ER, thereby forming new ER exit sites (ERES) containing the Sec24D coatomer (Sec24D-ERES). Based on this rationale, we hypothesize that the cellular process of cytisine-mediated chaperoning would increase neuronal resilience against pathogenic insults such as hyperoxidative stress by reducing the overall intensity of an apoptotic ER stress response.^8^ Although this is a feasible molecular mechanism to explain cytisine-mediated neuroprotection, a major question is whether or not the formation of new Sec24D-ERES due to upregulation of nAChRs in SNc DA neurons is sufficient for mediating neuroprotection in female parkinsonian mice. To address this critical question with clinically relevant implications for neuroprotection against Parkinson’s disease, we develop a novel transgenic mouse line with genetic mutations in the β2 subunit that cause β2* nAChRs to constitutively upregulate *in vivo* without the need for nicotinic ligands. Our newly created transgenic mouse line (called β2-upregulated mice) contains mutations in the M3-M4 intracellular loop of the β2 nAChR subunit that enhance β2* nAChR export from the ER, as well as prevent the ER retention of β2* nAChRs.^11^ Using these novel transgenic mice with constitutive genetically encoded β2* nAChR upregulation, for the first time, we show that only female mice demonstrate an upregulation of β2* nAChRs in SNc DA neurons, which causes neuroprotection in 6-OHDA-exposed female parkinsonian mice, and reduced apoptotic ER stress within SNc DA neurons.

More broadly, results from our study provide a novel and translationally relevant disease-modifying treatment avenue that could utilize combination drugs consisting of nicotinic ligands along with non-feminizing estrogen analogs for Parkinson’s disease in both male and post-menopausal female patients. In addition to developing important inroads into disease-modifying therapies for Parkinson’s disease, we envision that the β2-upregulated transgenic mice generated in this study will enable an understanding of the specific role of nAChR upregulation in other major neurological disorders such as addiction, anxiety, depression and dementia.^12^

## Materials and methods

### Mice

All experiments were conducted in accordance with Texas A&M University IACUC regulations and protocols (protocol # IACUC 2022-0252). ICR (CD1) and C57BL/6N mice were bred in house at the Texas A&M Institute for Genomic Medicine (TIGM, College Station, TX) and used to generate all β2-upgulated heterozygous (Het), homozygous (Hom) and wild-type (Wt) littermate mice for experiments. Mice were housed with littermates in ventilated cages and maintained on a 12-hour light-dark cycle, and food and water were provided *ad libitum*. All survival surgeries were conducted with continuous isoflurane anesthesia. Pain and suffering were minimized in 6-OHDA-treated animals by providing moistened peanut butter and Nutrical gel supplement, placed on the cage floor for easy access and to prevent weight loss following 6-OHDA lesions.

### Generation of Transgenic Mice

Single guide RNAs (sgRNAs) were designed using a proprietary algorithm from the CRISPR Core Partnership Program at Sigma Millipore (Merck KGaA, Darmstadt, Germany) and validated using the CEL I mutation detection assay (Sigma Millipore). The guide with the highest activity was selected for each target site and used to create single-stranded DNA oligonucleotide donors (ssDNA) synthesized by Sigma Millipore. Before injection, Cas9 mRNA (Sigma, cat #CAS9MRNA), sgRNAs, and ssDNAs were combined in TE buffer and centrifuged at 14,000 rpm for 10 minutes. The validated gRNA targeting sequence for the Chrnb2 gene was 5’ GCGCCAGCGGGAACGTGAG 3’. Donor oligonucleotides were designed to introduce specific single nucleotide polymorphisms (SNPs) at targeted locations to block gRNA rebinding and to maintain coding sequence integrity. SNPs were introduced at C>A, this conversion caused the amino acid (a.a) at 349 to change from leucine to methionine. SNPs introduced at AG>GC and two CG>GC sites, resulted in the conversion of arginine (R) to alanine (A) at a.a positions 365, 366, and 368. Additionally, a restriction enzyme recognition site for AccBSI was engineered within the oligonucleotide sequence to enable validation of successful edits. The donor oligo sequence retained the CRISPR gRNA target sequence to ensure accurate alignment during homology-directed repair (HDR) and included left and right homology arms flanking the target site to facilitate HDR. These features were specifically designed to ensure precise editing, allow functional validation through restriction site analysis, and support efficient HDR-based repair of the target locus.

Female C57BL/6N mice (3-5 weeks old) were superovulated by intraperitoneal (i.p.) injections of 5 IU Pregnant Mare Serum Gonadotropin (PMSG, ProSpec-Tany TechnoGene) followed 48 hours later by 5 IU Human Chorionic Gonadotropin (HCG, Sigma-Aldrich). Superovulated females were mated with 10-30-week-old males, and zygotes were harvested from the oviducts of euthanized females the next day in microinjection medium (HEPES-buffered DMEM with 5% FBS, ThermoFisher Scientific). Microinjections were performed under an inverted microscope equipped with a micromanipulator and Eppendorf Transjector 5246, using in house-pulled glass capillaries. Fertilized oocytes were injected into either the pronuclei or cytoplasm with CRISPR/Cas9 reagents, delivered via air-regulated flow at 90-115 psi to create a continuous flow of reagents. Pronuclear injections resulted in the distribution of reagents into both the pronuclei and cytoplasm due to continuous flow. Nine zygotes were transferred into each oviduct of pseudo-pregnant ICR (CD1) recipient females or cultured overnight at 37°C and transferred at the 2-cell stage.

Mice were genotyped between 12 and 17 days of age, prior to weaning. Primers were designed to specifically target the mutations and amplify regions surrounding the mutation sites. PCR amplification was conducted using LongAmp™ Taq Master Mix (New England Biolabs) under standard conditions. PCR products were purified using a PCR Clean-Up System kit, following the manufacturer’s protocol. Sanger sequencing was performed at the Texas A&M Sequencing Core to verify the mutation integrity and detect any unintended modifications within the cleavage site region. The primer sequences used for the wild-type PCR-based genotyping were as follows: Forward (Chrnb2 F) - 5’ CGTCACTAGCGTGTGTGTGC 3’ and Reverse (Chrnb2 Rw) - 5’ TCACGTTCCCGCTGGCGCCT 3’, which amplified a 178 bp region. For PCR-based genotyping in transgenic mice, the sequences used were as follows: Forward (Chrnb2 Fm) - 5’ CGCTTGCGAGCGGCCCAGGC 3’ and Reverse (Chrnb2 R) - 5’ CCTTGGTACTCACACTCTGGT 3’, which amplified a 265 bp region.

### Stereotaxic Surgeries

Surgical procedures were conducted as previously described.^8,9,13,14^ All surgeries were performed on three-month-old female mice under general anesthesia using isoflurane, with induction at 5% and maintenance at 1-2%, dispensed through a SomnoSuite Low Flow Anesthesia System (Kent Scientific, Torrington, CT). We used adeno-associated viruses (AAVs) to express proteins for visualization of acetylcholine (ACh)-evoked Ca^2+^ events in SNc DA neuron dendrites and dopamine release in the DLS. To specifically image Ca^2+^ activity in dendritic compartments of DA neurons, 1 μl of AAV9-TH-mCherry (10^13^ genome copies/ml; Vector Builder, cat # AAV9MP(VB220210-1216jhh)-C) and 1 μl of AAV1-Syn-GCaMP6f (10^13^ genome copies/ml; Addgene viral prep #100837-AAV1) were mixed and co-injected into the SNc using a beveled glass injection pipette at a flow rate of 0.5 nl/min attached to a QSI Dual Microliter Syringe Pump (Stoelting, cat # 53325). Coordinates for SNc surgeries were 3.0 mm posterior to bregma, 1.5 mm lateral to midline, and 4.2 mm ventral to the pial surface. The injection pipette was left in place for 5-10 minutes after injection, then gradually withdrawn. Surgical wounds were closed with tissue adhesive (Amazon). Mice were sacrificed three weeks later for acute brain slice imaging. To image dopamine release, 1 μl of AAV9-hsyn-GRAB_DA2m(33087905) (10^13^ genome copies/ml; Addgene viral prep #140553-AAV9, RRID: 140553) was injected into the DLS at a flow rate of 0.5 nl/min. Coordinates for stereotaxic injections into the dorsolateral striatum (DLS) were 0.8 mm anterior to bregma, 2.0 mm lateral to midline, and 2.4 mm ventral to the pial surface. Mice were sacrificed four weeks later for acute brain slice imaging. For unilateral striatal 6-OHDA lesions, 5 mg/ml stock solutions of 6-OHDA (Sigma, St. Louis, MO) were prepared in 0.9% saline with 0.2% ascorbic acid and frozen at -80°C until use. Mice received a unilateral stereotaxic injection of 2 μl of stock solution (10 μg of 6-OHDA in total) into the DLS at a rate of 1 μl/min, using the same DLS coordinates described above. Following the lesion, all mice were monitored daily for significant weight loss before proceeding with behavioral testing.

### Imaging of acute mouse brain slices

Acute mouse brain slices for imaging were obtained as previously described.^13–15^ Briefly, coronal brain slices (300 µm thick) of the DLS or SNc were cut using a Microslicer 01 N (Ted Pella) in a slicing solution consisting of (in mM): 194 sucrose, 30 NaCl, 4.5 KCl, 1.2 NaH_2_PO_4_, 26 NaHCO_3_, 10 D-glucose, and 1 MgCl_2_ and saturated with 95% O_2_ and 5% CO_2_ (pH 7.2). Slices were incubated and recorded in artificial cerebrospinal fluid (aCSF) comprising (in mM): 126 NaCl, 2.5 KCl, 1.24 NaH_2_PO_4_, 26 NaHCO_3_, 10 D-glucose, 2.4 CaCl_2_, and 1.3 MgCl_2_ saturated with 95% O_2_ and 5% CO_2_ (pH 7.4) and 500 nM of the acetylcholinesterase inhibitor, Donepezil (Sigma, cat # D6821). Slices were incubated in a solution of 25% slicing solution and 75% recording solution at 34°C for 25 minutes, then maintained at room temperature (RT) for the duration of the experiment.

Imaging was performed as previously described.^13–15^ Slices were imaged using an Olympus FV3000 upright laser-scanning confocal microscope with a 40x water immersion objective lens, numerical aperture (NA) of 0.8. 488 and 561 nm LED-based excitation wavelengths were used for obtaining images. For DLS imaging, the 488 nm line intensity was set at 10% maximum output to visualize GRABDA fluorescence and recordings were obtained with 1x digital zoom. For SNc imaging, the 488 nm and 561 nm LED intensity was set at 10% to visualize GCaMP6f and TH fluorescence respectively and recordings were obtained at 3x digital zoom. Confocal parameters (high voltage, gain, offset, laser power, and aperture diameter) were held constant for GRABDA and GCaMP6f imaging sessions. Events were recorded at 1 frame per second (FPS) for 600 seconds.

All drugs were bath perfused using a peristaltic pump (Harvard Apparatus), and the bath perfusion timing was set prior to each imaging session. To assess changes in dopamine release in the DLS or Ca^2+^ flux in TH+ dendritic compartments, spontaneous activity was recorded for the first 100 seconds, followed by the bath application of 300 µM ACh (Sigma, cat# A2661-25G) for the remaining 500 seconds. To determine the contribution of β2* nAChRs to the observed response, 1 µM of dihydro-β-erythroidine (DhβE, Tocris, cat # 2349) was perfused into the bath and slices were incubated with DhβE for 5 minutes prior to recording. Activity with DhβE alone was recorded for the first 100 seconds, followed by the bath application DhβE and ACh for the remaining 500 seconds.

### Apomorphine Rotations

Apomorphine-induced rotational behavior was performed as previously described.^8,9^ Mice were acclimated individually in 5-gallon buckets for 15 minutes. Following acclimation, mice received an intraperitoneal (i.p) injection of 0.5 mg/kg apomorphine (Sigma, cat# 1041008) dissolved in 0.9% saline. Rotational behavior was recorded for 15 minutes to obtain the number of contralateral rotations for each mouse and Ethovision XT (Noldus, Leesburg, VA) was used to analyze contralateral rotations with arena diameters, trial settings, and detection settings kept constant for all mice across all behavioral testing sessions.

### Immunostaining

Mice were deeply anesthetized with isoflurane, then transcardially perfused with PBS, immediately followed by 10% formalin (VWR, cat# 100496-506). After extraction, brains were preserved in 10% formalin at 4°C for 48 hours, then dehydrated in 30% sucrose in PBS (Sigma, cat# S7903) for 48 h. All brain sections (40 μm thickness) were prepared using a microtome (Leica) and stored in 0.01% sodium azide in PBS (Sigma, cat # S2002). To maintain precise rostrocaudal orientation, midbrain sections were sequentially collected in 96-well plates. For immunostaining, sections were washed twice in PBS for 5 minutes each, then permeabilized and blocked with 0.5% Triton X-100 (Sigma, cat # X100) + 10% normal goat serum (NGS) (Abcam, cat # ab7481, RRID: AB_2716553) for 45 minutes at RT with constant agitation. Sections were then labeled with the appropriate primary antibodies in 0.05% Triton X-100 + 1% NGS in PBS at 4°C overnight. The next day, sections were incubated with the appropriate secondary antibodies in 0.05% Triton X-100 + 1% NGS in PBS for 1 h at RT. Sections were mounted on microscope slides using PBS and protected with fluoromount (Diagnostic Biosystems, cat # K024) for imaging. Slides were stored at 4°C until imaging. Primary antibodies were chicken anti-TH (1:2000; Abcam, cat # ab76442, RRID: AB_1524535), rabbit anti-C/EBP homologous protein (CHOP) (1:1000; Cell Signaling, cat # 5554, RRID: AB_10694399), rabbit anti-Sec24D (1:500; ThermoFisher, cat # PA5-65793, RRID: AB_2662256), and mouse anti-glial fibrillary acid protein (GFAP) (1:2000; Invitrogen cat # 14-9892-82, RRID: AB_10598206). Secondary antibodies were goat anti-chicken Alexa Fluor 594 (1:2000; Abcam, cat # ab150176, RRID: AB_2716250), goat anti-rabbit Alexa Fluor 488 (1:2000 for CHOP, 1:1000 for Sec24D; Abcam, cat # ab150077, RRID: AB 2630356), and goat anti-mouse Alexa Fluor 647 (1:2000, Abcam, cat # ab150115, RRID: AB_2687948).

### Quantification of Sec24D-ERES number in SNc DA neurons

Midbrain sections were immunostained for TH (goat anti-chicken Alexa Fluor 594 secondary antibody) and Sec24D-ERES (goat anti-rabbit Alexa Fluor 488 secondary antibody) and imaged using an inverted Olympus FV3000 laser-scanning confocal microscope equipped with a 60x oil immersion objective lens (NA 1.35). LED excitation wavelengths at 488 nm for Sec24D-ERES and 561 nm for TH fluorescence were used. Microscope settings, including LED power, high voltage (HV), gain, offset, and aperture diameter were optimized to prevent image saturation and kept consistent across all imaging sessions. Individual TH+ cell bodies of SNc DA neurons in midbrain sections were imaged at 60x magnification with 3x digital zoom. Each SNc DA neuron was first imaged using TH to focus on an optical plane where the nucleus was clearly defined and visible as a dark halo. Four optical sections were then imaged for TH and Sec24D-ERES on either side of this central reference optical plane for a total of 8 optical sections per neuron with a step size of 0.47 μm. Images were obtained for randomly selected cells from ∼8 midbrain sections in each mouse, resulting in images of 70-170 individual SNc DA neurons per genotype per mouse.

Quantification of the number of Sec24D-ERES structures per neuron was performed using ImageJ software by a blinded experimenter as follows: A maximum intensity z-projection of Sec24D-ERES was generated for each TH+ SNc DA neuron, and Sec24D-ERES structures in each neuron from the z-projected image was manually thresholded. Regions of interest (ROIs) for Sec24D-ERES were defined using the manually thresholded images and Analyze Particles feature in ImageJ, with an ROI area constraint of 0.01-1000 μm². ROIs were used to determine the number of ERES per TH+ neuron in the SNc. In order to address sampling bias, the number of ERES were plotted per cell, then averaged per section, and finally averaged per mouse.

### Quantification of ACh-evoked Ca^2+^ responses in dendrites of SNc DA neurons

As described above, we utilized AAV-mediated co-expression of TH-mCherry and hSyn-GCaMP6f in the mouse SNc to specifically visualize and image ACh-evoked Ca^2+^ responses in SNc DA neuron dendrites from live mouse midbrain slices. XY time-series movies were drift-corrected along the x-y axis using the Turboreg plugin in ImageJ. For each analyzed movie, a corresponding z-stack image of mCherry and GCaMP6f expression was used to generate red and green co-localized maximum intensity projection images. Co-localization of mCherry and GCaMP6f fluorescence was used to identify Ca^2+^ responses within TH+ dendritic compartments of SNc DA neurons.

Regions of interest (ROIs) were manually drawn on TH+ dendritic structures, with a consistent ROI size of 2.64 μm, maintained across all slices. ROIs displaying an increase in ACh-evoked Ca^2+^ responses were selected for analysis. Identical ROIs were used to analyze Ca^2+^ responses with ACh only as well as ACh-evoked Ca^2+^ responses in the presence of DhβE. To quantify changes in fluorescence intensity over time, area under the curve (AUC) measurements were obtained using Origin 2019 v9.6 (OriginLab). Mean gray values of GCaMP6f fluorescence intensity were plotted as line graphs, and the integrate tool in the gadgets menu of Origin 2019 was applied to each trace. Baseline AUC adjustments were made to account for photobleaching. The AUC was measured over 470 seconds for all traces, providing a standardized approach for comparing ACh-evoked Ca^2+^ flux across experiments. To calculate the percentage inhibited by DhβE, the AUC from the trace with ACh-evoked Ca^2+^ responses in the presence of DhβE was divided by the AUC from the corresponding ROI trace with Ca^2+^ response with ACh only. This yielded the percentage of response not inhibited by DhβE. The percentage of DhβE inhibition of ACh-evoked response was then calculated as 100 minus percentage of ACh-evoked Ca^2+^ response not inhibited by DhβE.

### Quantification of ACh-evoked dopamine release in the DLS

The analysis of ACh-evoked dopamine release was conducted using the same imaging and drift correction methods as those applied for ACh-evoked Ca^2+^ flux. In this case, ROIs were manually drawn to encompass the entire field of view (FOV) and used to extract mean gray values of GRABDA fluorescence intensities. Only slices showing an increase in ACh-evoked DA release were included in the analysis. AUC for GRABDA traces and percentage of dopamine release inhibition by DhβE was calculated using the same method as described above for ACh-evoked Ca^2+^ responses in the SNc.

### Quantification of SNc DA neuron loss

SNc DA neurodegeneration following 6-OHDA lesion was quantified by comparing ratios of area of TH fluorescence between the unlesioned and lesioned SNc. For each mouse, 10 midbrain sections (spaced 120 µm apart) were collected, and 8 sections in which the SNc and VTA were visually distinct were selected for analysis. Sections were imaged using an Olympus VS120 virtual slide scanning microscope with a 10x air objective lens. TH labeled DA neurons and processes in the SNc were imaged with 561 nm excitation and an exposure time of 400 ms per section. Rectangular ROIs (10 mm²) were drawn on both the lesioned and unlesioned SNc within each image. A 20 μm thick optical z-stack was acquired for every section under consistent imaging parameters. Image analysis was performed in Fiji, where a sum z-projection was generated and the polygon tool used to manually outline the ROIs corresponding to the lesioned and unlesioned SNc. Thresholding of TH labeled neurons and processes was applied to isolate TH+ cell bodies and processes for subsequent quantification of area of TH labeled structures for each section. The ratios of TH+ area between the lesioned and unlesioned SNc were compared by obtaining ratios from summed areas of 8 sections per mouse. Area ratios were used for each section to minimize variability introduced by immunostaining artefacts across sections and mice.

### Subcellular quantification of CHOP localization in SNc DA neurons

To quantify the extent of nuclear translocation of CHOP, SNc DA neurons from midbrain sections on the 6-OHDA lesioned side were imaged using an inverted Olympus FV3000 laser-scanning confocal microscope. The 488 nm LED wavelength was used to image CHOP, and the 594 nm wavelength was used to image TH in the SNc. TH+ cell bodies were imaged at 60x magnification with a 4x digital zoom in lesioned SNc brain sections. Single-plane images were collected from 36-39 TH+ neurons per genotype. Planes were selected only when the nucleus was clearly visible as a dark halo in the TH imaging channel. The lack of TH labeling within the nucleus allowed for distinct identification of nuclear and cytosolic compartments. Image analysis was performed using Fiji software. ROIs were manually delineated using the polygon tool to encompass the nucleus and the whole cell boundary for each TH+ neuron, which enabled quantification of CHOP fluorescence intensity in both regions. Additionally, the percentage of CHOP nuclear translocation was measured by obtaining the ratio of nuclear to whole cell CHOP fluorescence intensity.

### Quantification of glial fibrillary acid protein (GFAP) expression in the SNc

Astrocyte reactivity following the 6-OHDA lesion was assessed using GFAP labeling as a marker of reactivity in three mouse midbrain sections per mouse where the SNc and VTA were visually distinct. Sections were imaged with an Olympus VS120 virtual slide scanning microscope equipped with a 10x air objective lens. GFAP fluorescence was imaged with 647 nm excitation and a 200 ms exposure time, and TH fluorescence was imaged with 561 nm excitation and a 30 ms exposure time. Rectangular ROIs of 10 mm² were drawn to include TH+ structures on both the lesioned and unlesioned sides of the SNc in a single image. A 20 μm thick optical z-stack was acquired for each section in extended focus image (EFI) z-mode, selecting images with the highest sharpness values. Imaging parameters were kept consistent across sections and mice. Image analysis was performed using Fiji software. The polygon tool was used to manually delineate ROIs for the lesioned and unlesioned SNc, and GFAP-integrated fluorescence intensity was measured within these ROIs. Integrated intensities from the three sections were averaged per mouse.

### Statistical Analysis

All statistical analyses were performed using Origin Lab. For the apomorphine behavioral assay, a two-way repeated measures analysis of variance (ANOVA) was used to examine the effects of genotype on rotational behavior with genotype and time as the two variables. The day of testing served as a repeated measure and genotype as the between-subject factor. *p* < 0.05 with a high F statistic was considered statistically significant. For all other datasets in this study, data for each condition were first tested for normality using a Shapiro-Wilk test. A two-sample t-test was used for normally distributed data, while a Mann-Whitney test was used for non-normally distributed data and *p* < 0.05 was considered as statistically significant. Figure legends contain sample sizes and the specific statistical tests used for each experiment.

## Results

### Generation of transgenic β2-upregulated mice with enhanced ER export of β2* nAChRs

We previously showed that mutating leucine (L) to methionine (M) at amino acid (a.a) position 346 in the M3-M4 intracellular loop of the β2 nAChR subunit results in the formation of an LFM motif (a.a 344-346) which enhances the ER export of β2* nAChRs.^11^ This study also showed that mutating arginine (R) in the RRQR ER retention motif to alanine (A), AAQA (a.a 365-368) in the M3-M4 intracellular loop of β2 nAChR subunits functionally disrupts ER retention β2* nAChRs (Fig. 1A).^11^ Thus, together, the two mutations (344-LFM-346 and 365-AAQA-368) in the M3-M4 intracellular loop of β2 nAChR subunits act synergistically to increase pentameric nAChR exit from the ER, resulting in a two-fold upregulation of β2* nAChRs at the cellular plasma membrane.^11^ Based on these previous findings, we used CRISPR-Cas9 gene editing to generate a novel transgenic mouse model, called β2-upregulated mice, with both mutations (344-LFM-346 and 365-AAQA-368) in the M3-M4 loop of β2 nAChR subunits that would together upregulate β2* nAChRs *in vivo* (Fig. 1B). For all mice described in this study, these two genetic mutations in the M3-M4 loop of the mouse *Chrnb2* gene were confirmed using PCR-based genotyping. Wild-type (Wt) and mutation-specific primer sequences, as described in methods, were employed to differentiate between Wt, Het and Hom genotypes. Each mouse was screened with both Wt and mutant primer pairs, with a band at 178 bp indicating the Wt allele and a band at 265 bp indicating the mutant allele. A single band at 178 bp with only the Wt primer identified the mouse as Wt, while a band at 265 bp with the mutant primer pair and a 178 bp band with the Wt primer pair identified a Het mouse. Finally, a 265 bp band with the mutant primer pair, and no band with the Wt primer pair was used to identify mice as Hom (Fig. 1C). Over the course of this study, all β2-upregulated transgenic mice bred normally and did not display any overt behavioral or physiological abnormalities.

**Figure 1.**
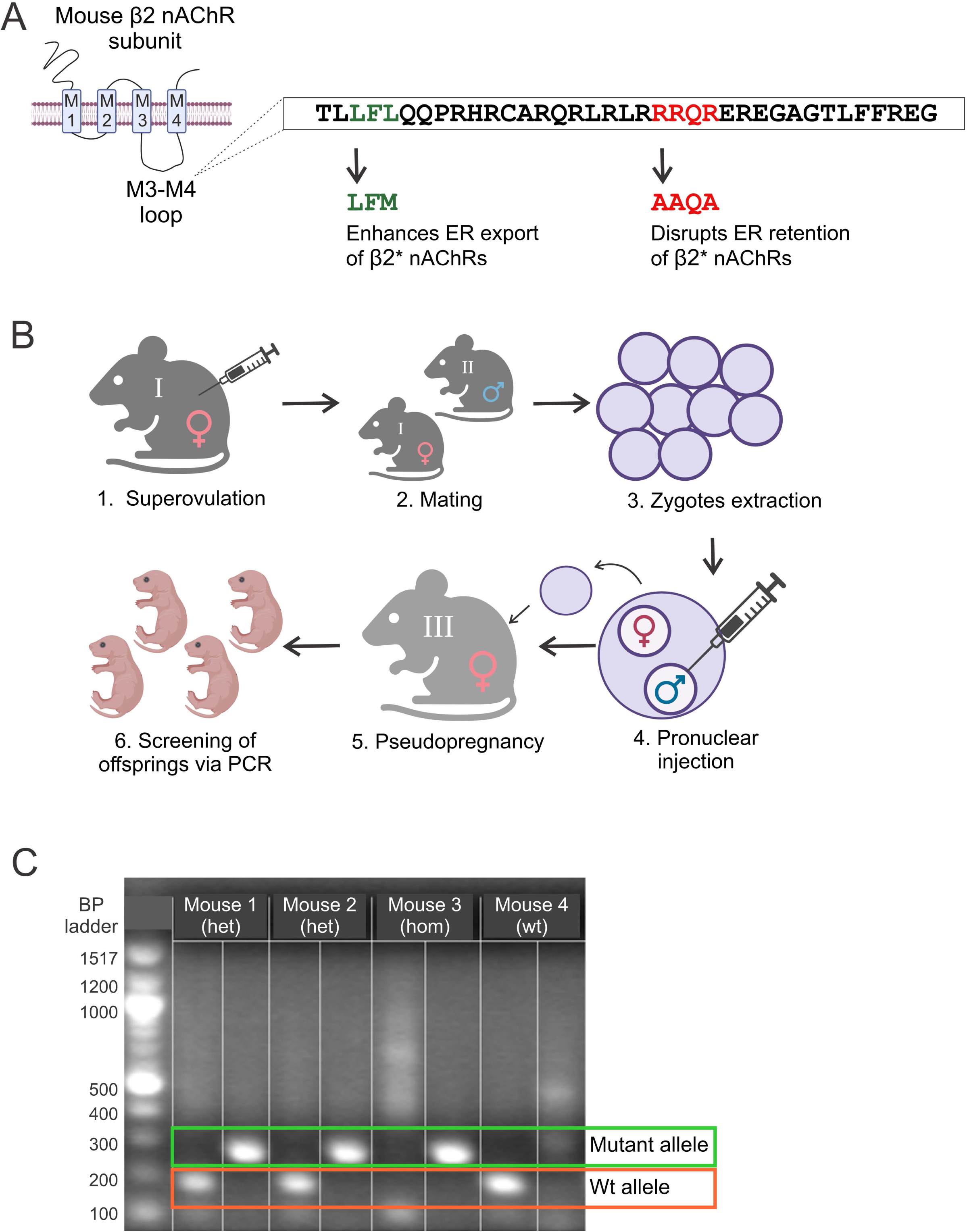
Creation and genotyping of β2-upregulated transgenic mice designed to mimic nAChR upregulation. **(A)** Schematic of the M3-M4 intracellular loop of mouse β2 nAChR subunits, illustrating the two-point mutations engineered to generate β2-upregulated mice. The first mutation (LFL→LFM) enhances ER export, whereas the second (RRQR→AAQA) disrupts the ER retention signal. **(B)** Stepwise illustration of the process to generate β2-upregulated mice using CRISPR/Cas9 technology (a detailed description is provided in the Materials and Methods section). **(C)** Agarose gel electrophoresis showing PCR amplification products. DNA samples from four different mice are shown, indicating the presence of wild-type (Wt) and mutant alleles. Each mouse was screened using a Wt primer (first column) and a mutant primer (second column). A 178-bp band (outlined in orange) confirmed the Wt allele, whereas a 265-bp band indicated the mutant allele. Panels A and B were created using BioRender.

### Only female β2-upregulated transgenic mice demonstrate an increase in Sec24D-containing ERES in SNc DA neurons

LFM motifs have been shown to specifically bind to the Sec24D isoform,^16^ a core component of ERES that nucleates the formation of ERES and aids in the exit of captured cargo from cellular ER. Because we introduced a new LFM motif in the M3-M4 loop of β2 nAChR subunit of β2-upregulated transgenic mice (Fig. 1C), we asked whether female and male Het and Hom β2-upregulated mice demonstrate an increase in Sec24D-containing ERES due to enhanced β2* nAChR export out of the ER in SNc DA neurons. To quantify Sec24D-ERES structures in TH+ DA neurons, we co-immunostained midbrain sections with Sec24D and TH (Fig. 2A) and quantified the number of Sec24D-ERES in SNc DA neurons at the cellular level. To control for sampling bias, these data were averaged for multiple cells within each section, as well as across multiple sections for each mouse (Fig. 2B). Data for Het and Hom female and male mice were pooled together within female and male sexes because no significant differences were found between Het and Hom mice. We found that both Het and Hom β2-upregulated transgenic female mice exhibited a significant increase in the number of Sec24D-ERES when compared to Wt littermate female mice (Number of ERES per cell: Wt, 22.43 ± 1.24; β2-upregulated Het and Hom, 33.03 ± 1.31, *p* < 0.001; Number of ERES per section: Wt: 22.54 ± 2.03; β2-upregulated Het and Hom, 33.19 ± 1.96, *p* = 1.63 x 10⁻⁷ ; Number of ERES per mouse: and Wt, 22.36 ± 3.64, β2-upregulated Het and Hom, 32.96 ± 2.77, *p* = 0.04; Mann-Whitney test) (Fig. 2A and 2B). By contrast, in male mice, no significant differences were observed between Wt male littermates and β2-upregulated male transgenic mice (Number of ERES per cell: Wt, 28.44 ± 1.89; β2-upregulated Het and Hom, 28.03 ± 1.28, *p* = 0.8; Number of ERES per section: Wt, 29.44 ± 3.22; β2-upregulated Het and Hom, 28.63 ± 2.1; *p* = 0.71; Number of ERES per mouse: Wt, 31.18 ± 6.05; β2-upregulated Het and Hom, 29.86 ± 3.71, *p* = 0.73; Mann-Whitney test) (Fig. 2A and 2B). Taken together, these results show a novel sex-specific effect of β2* M3-M4 loop nAChR mutations, whereby there is enhanced export of functional β2* nAChRs from the ER of SNc DA neurons in female but not in male β2-upregulated transgenic mice. The observed sex difference in Sec24D-ERES upregulation motivated further assessment of whether β2* nAChRs are functionally upregulated in SNc DA neurons of female transgenic mice.

**Figure 2.**
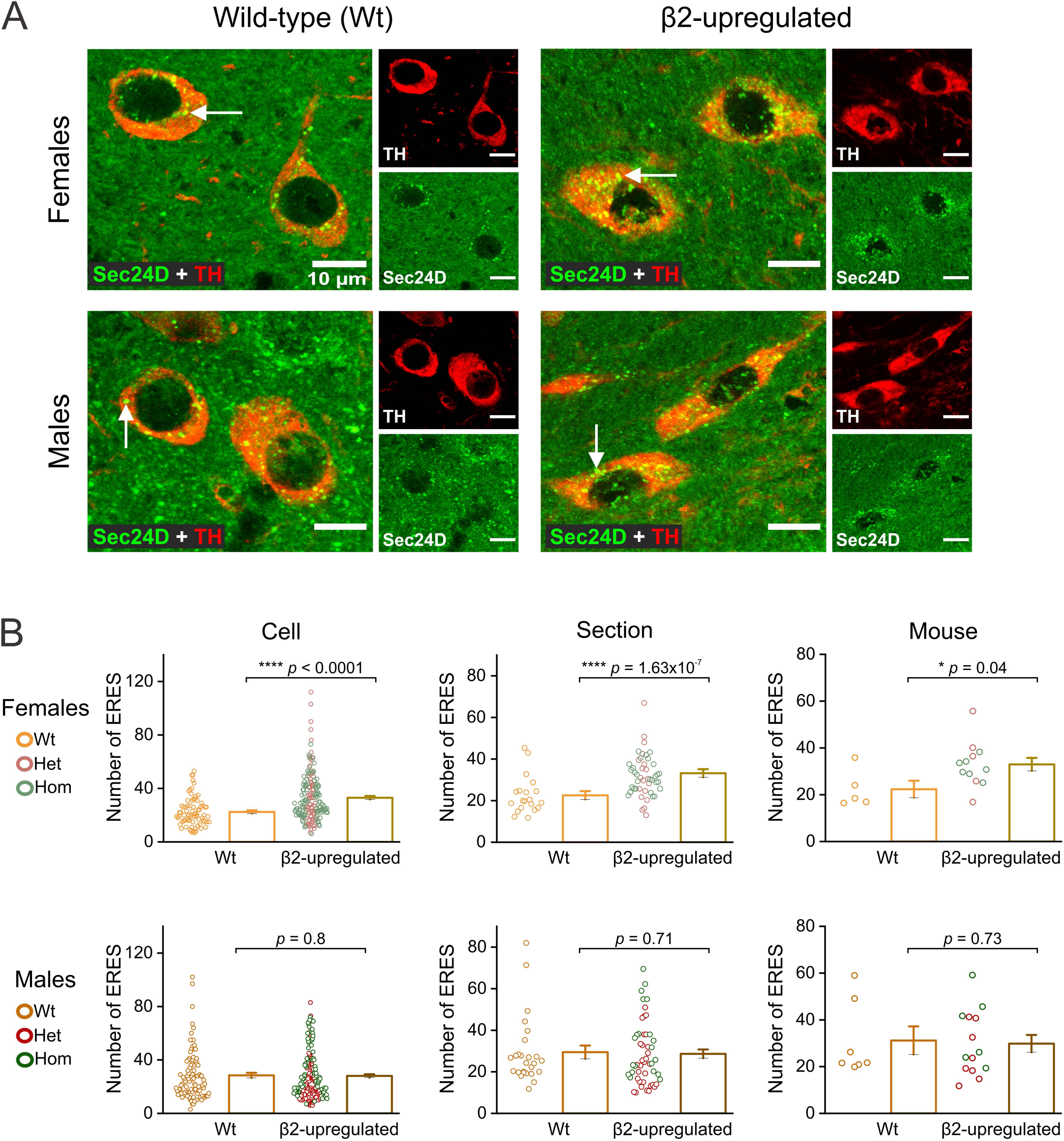
Sec24D-containing ERES are upregulated only in female β2-upregulated female mice. **(A)** Representative confocal images of SNc DA neurons from Wt and β2-upregulated female (top) and male (bottom) mice stained for endogenous TH (red) and Sec24D-ERES (green). Merged images and separated channels are shown for TH (red) and Sec24D (green) staining. White arrows point to Sec24D-ERES puncta in TH+ SNc DA neurons. Scale bar, 10 µm. **(B)** Bar graphs quantifying number of Sec24D-ERES puncta per cell (left), per section (middle), and per mouse (right) for female and male Wt and transgenic mice. Cell sampling: Female Wt n=75; Female β2-upregulated n=177 (Heterozygous (Het) n=73, Homozygous (Hom) n=104); Male Wt n=95; Male β2-upregulated n=188 (Het n=111, Hom n=77). Section sampling (4 sections per mouse): Female Wt n=20; Female β2-upregulated n=48 (Het n=20, Hom n=28); Male Wt n=26; Male β2-upregulated n=51 (Het n=30, Hom n=21). Mouse sampling: Female Wt n=5; Female β2-upregulated n=12 (Het n=5, Hom n=7); Male Wt n=7; Male β2-upregulated n=14 (Het n=8, Hom n=6). All p-values are based on Mann-Whitney tests and error bars represent ± S.E.M.

### β2* nAChRs are functionally upregulated in SNc DA neuron dendrites of female β2-upregulated transgenic mice

Based on our finding that the number of Sec24D-ERES is increased in SNc DA neurons of female but not male β2-upregulated mice (Fig. 2A and B), we next sought to assess whether β2* nAChRs are functionally upregulated in the SNc DA neurons of female β2-upregulated transgenic mice. Since previous reports have shown that β2* nAChRs can traffic to neuronal dendrites and couple with other Ca^2+^ channels at dendritic locations,^17–20^ we hypothesized that β2* nAChR activation by ACh would cause a local depolarization in the dendrites of SNc DA neurons, resulting in quantifiable increases of Ca^2+^ influx via L-type voltage-gated calcium channels (L-type VGCCs). We further rationalized that Ca^2+^ influx into dendritic ROIs would be inhibited by the β2* nAChR-specific antagonist, DhβE.^21^ Based on this rationale, we performed experiments to assess functional β2* nAChR upregulation in β2-upregulated transgenic mice using bath ACh-evoked Ca^2+^ responses in SNc DA neuron dendrites with or without the β2* nAChR-specific antagonist, DhβE. To do this, we stereotaxically co-injected the SNc of female Wt, Het and Hom mice with two AAVs: One AAV expressing GCaMP6f driven by the neuron-specific hSyn1 promoter, and a second AAV expressing mCherry, driven by a TH minimal promoter. AAV-mediated co-expression of GCaMP6f and mCherry enabled specific visualization of Ca^2+^ fluxes in the dendritic compartment of SNc DA neurons. In both Wt and β2-upregulated female mice, bath application of 300 µM ACh to midbrain slices resulted in a robust increase of GCaMP6f fluorescence in the TH-mCherry labeled dendrites of SNc DA neurons (Fig. 3A and 3C). To specifically assess the contribution of β2* nAChRs to dendritic Ca^2+^ influx responses, we quantified Ca^2+^ responses in the same ROIs following bath co-application of 1 μM DhβE + 300 μM ACh. When compared to Wt female littermates, ACh-induced Ca^2+^ influx into SNc DA neuron dendritic ROIs showed an increase in the AUC which was reduced to a significantly greater degree by co-applied DhβE in β2-upregulated transgenic female mice than Wt mice (Fig. 3C) [Wt mice AUC (A.U. x sec), ACh only: 31245.27 ± 1992.56, DhβE + ACh: 24196.09 ± 1689.99, *p* = 1.77 x 10^-4^, two-sample t-test; β2-upregulated mice AUC (A.U. x sec), ACh only: 32963.22 ± 2018.46, DhβE + ACh: 14109.04 ± 720.97, *p* < 0.0001 two-sample t-test]. In addition, when compared to Wt female mice, the percentage of DhβE-sensitive ACh-evoked Ca^2+^ responses was ∼2-fold greater in β2-upregulated female mice when compared to Wt mice (Wt: 27 ± 0.02 %; β2-upregulated: 51 ± 0.02 %, *p* < 0.0001; two-sample t-test) (Fig. 3C). Taken together, these data show that female β2-upregulated transgenic mice demonstrate a constitutive genetically-encoded functional upregulation β2* nAChRs in the dendrites of SNc DA neurons.

**Figure 3.**
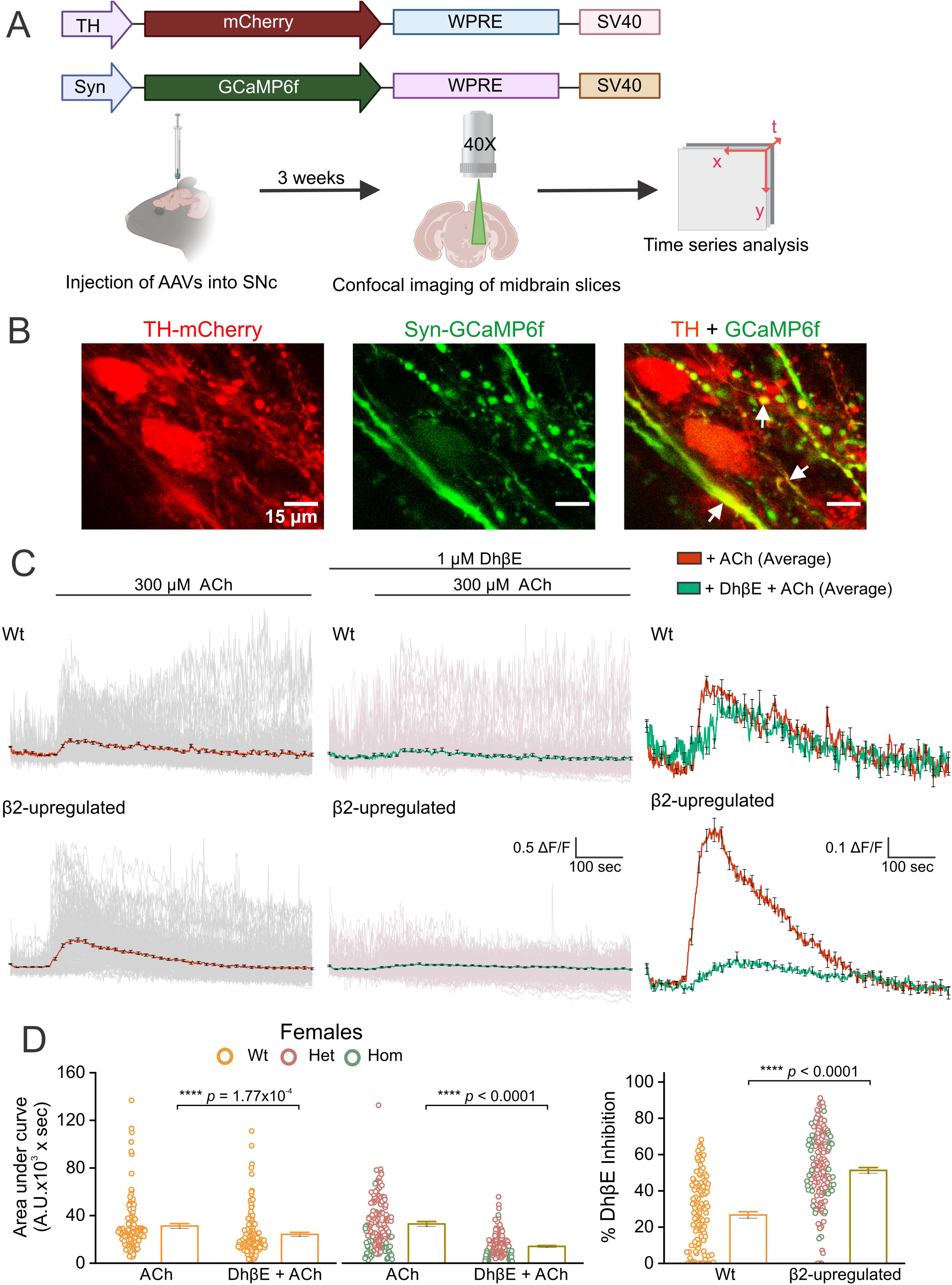
ACh-evoked DhβE-sensitive Ca^2+^ influx is increased in TH+ dendritic compartments in SNc of β2-upregulated female mice. (A) Schematic of AAV9-TH-mCherry and AAV1-Syn-GCaMP6f injected into the SNc of female mice. Three weeks later, confocal imaging was performed on acute midbrain slices, followed by time-series analysis to measure changes in GCaMP6f fluorescence intensity over time. **(B)** Representative time-compressed confocal images of TH-mCherry (red) and GCaMP6f (green) from β2-upregulated female mice. Scale bar, 15 µm. White arrows indicate TH+ dendritic compartments where Ca^2+^ influx was increased by bath application of ACh and inhibited by DhβE. **(C)** Traces of GCaMP6f fluorescence intensity in TH+ dendritic compartments from Wt (top) and β2-upregulated (bottom) mice during bath application of ACh (left, individual region of interest (ROI) traces in grey and average in red), co-application of DhβE + ACh (middle, individual ROI traces in pink and average in green), and averaged traces from all ROIs (right). **(D)** Bar graphs showing the area under the curve (AUC) per ROI with bath application of ACh and co-application of DhβE + ACh (left) and the percentage of DhβE inhibition (right). ROI sampling: Wt n = 123 and β2-upregulated n = 175 (Het n = 139, Hom n = 36). All p-values are from Mann-Whitney tests and error bars represent ± S.E.M. Panel A was created using BioRender.

### β2* nAChRs are functionally upregulated in the axonal terminals of SNc DA neurons within the DLS of female β2-upregulated transgenic mice

Having observed functional upregulation of β2* nAChRs in SNc DA neuron dendrites of β2-upregulated transgenic female mice, we next asked whether axon terminals of SNc DA neurons in the DLS of these transgenic mice also display functional upregulation of β2* nAChRs. Since previous studies have shown that ACh-evoked activation of nAChRs in DA axonal terminals within the DLS causes robust striatal dopamine release,^22–24^ we employed the optogenetic dopamine sensor, GRABDA^25^ to measure ACh-evoked dopamine release in live DLS slices as a functional assay for nAChR upregulation in axonal terminals. AAVs were used to express GRABDA in the DLS of female Wt and transgenic mice, and confocal imaging was performed in live striatal slices four weeks after AAV injection (Fig. 4A). For both Wt and β2-upregulated transgenic mice, bath application of ACh caused a robust increase in GRABDA fluorescence across entire FOVs, and this ACh-evoked GRABDA response was strongly inhibited in the presence of co-applied DhβE (Fig. 4B and C). To quantify GRABDA responses, we measured the AUC for each FOV with ACh and ACh + DhβE. In all cases, DhβE caused a nearly 4-fold inhibition of ACh-evoked increases in GRABDA fluorescence (Fig. 4D) [Wt mice AUC (A.U. x sec), ACh only: 9205.41 ± 1184.2, DhβE + ACh: 2566.46 ± 227.75, *p* = 0.002, two-sample t-test; β2-upregulated mice AUC (A.U. x sec), ACh only: 9503.28 ± 1241.98, DhβE + ACh: 1647.91 ± 144.81, *p* < 0.0001, two-sample t-test] (Figs. 4B, 4C). Importantly, the percentage of DhβE-induced inhibition of ACh-evoked GRABDA responses was ∼10% greater in β2-upregulated female mice compared to Wt (Wt: 71 ± 0.03 %; β2-upregulated: 81 ± 0.02 %, *p* < 0.005; two-sample t-test) (Fig. 4D). These findings show that β2-upregulated transgenic female mice do display a small but significant functional upregulation of β2* nAChRs in the striatal axonal terminals of SNc DA neurons.

**Figure 4.**
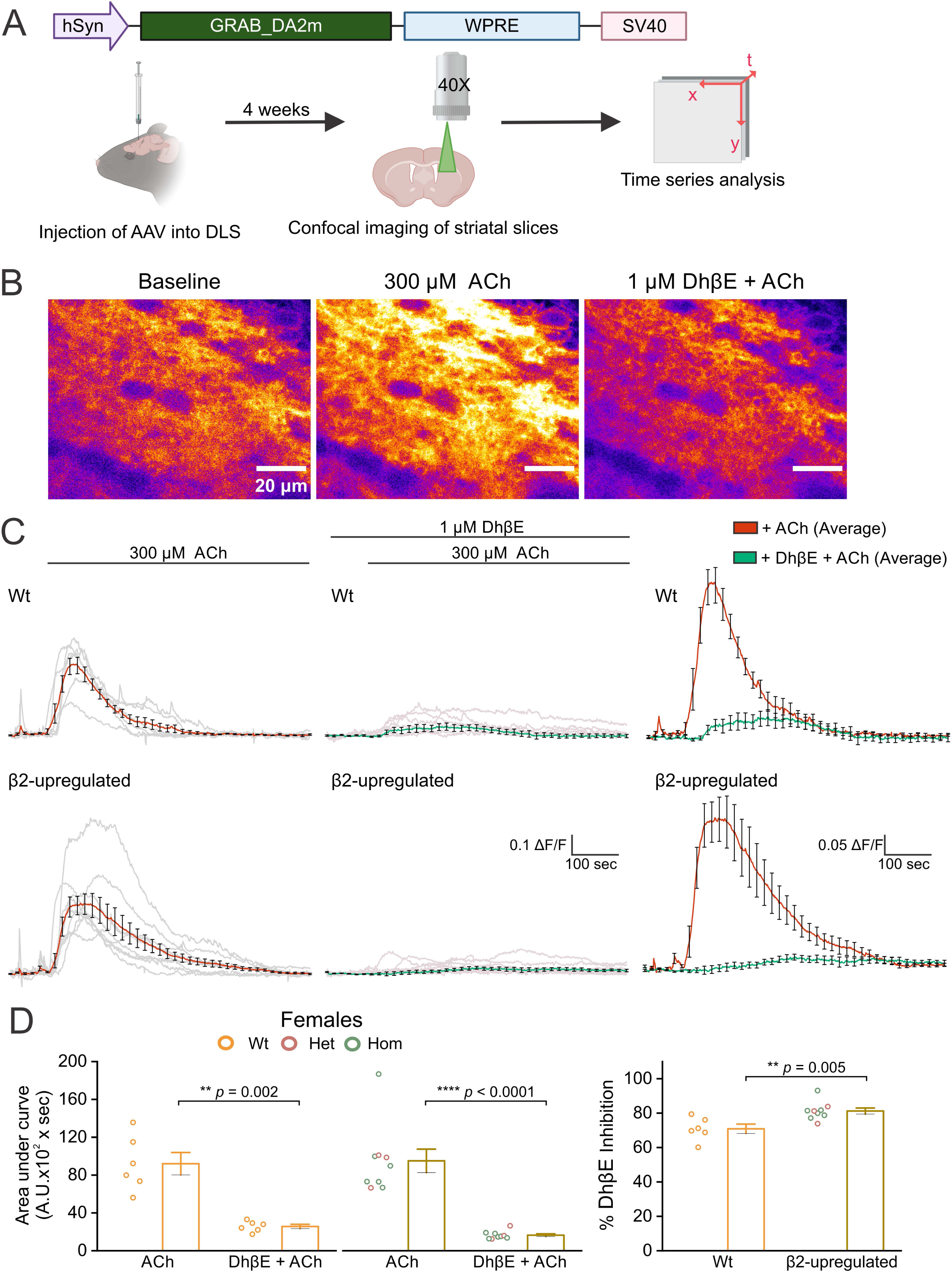
ACh-evoked DhβE-sensitive dopamine release is increased in the DLS of β2-upregulated female mice. **(A)** Schematic of AAV9-hsyn-GRAB_DA2m (GRABDA) injected into DLS of female mice. Four weeks later, confocal imaging was performed on acute mouse DLS brain slices, followed by time series analysis to measure changes in GRABDA fluorescence intensity over time. **(B)** Representative time-compressed confocal images of GRABDA at baseline (left), followed by bath application of ACh (middle), and co-application of DhβE + ACh (right). Scale bar, 20 µm. **(C)** Traces of GRABDA fluorescence intensity from Wt (top) and β2-upregulated (bottom) mice during bath application of ACh (left, traces per FOV in grey and average in red), co-application of DhβE + ACh (middle, traces per FOV are in light pink and average in green), and average traces from multiple FOVs (right). **(D)** Bar graphs showing the average area under the curve with bath application of ACh and co-application of DhβE + ACh (left) and the percentage of DhβE inhibition per mouse (right). n is 1-2 FOVs from 1-2 slices per mouse, averaged per mouse: Wt n = 6 and β2-upregulated n = 9 (Het n = 3, Hom n = 6). All p-values are from two-sample t tests and error bars represent ± S.E.M. Panel A was created using BioRender.

### Female 6-OHDA lesioned β2-upregulated mice display reduced apomorphine rotations

Our data thus far show that only female β2-upregulated transgenic mice display an increase in Sec24D-ERES which is utilized by β2* nAChRs to exit the ER (Fig. 2), and that β2* nAChRs in female transgenic mice are functionally upregulated within SNc DA neuron dendrites (Fig. 3) and in striatal axonal terminals of SNc DA neurons (Fig. 4). Based on these data and our previous studies showing that cytisine confers neuroprotection only in female mice, potentially via β2* nAChR upregulation, we next sought to determine if constitutive upregulation of β2* nAChRs in female transgenic mice without cytisine is neuroprotective in SNc DA neurons in a preclinical mouse model of 6-OHDA-induced parkinsonism. To assess neuroprotection in the β2-upregulated female transgenic mice, we unilaterally injected 10 μg of 6-OHDA into the DLS of mice to induce a unilateral loss of SNc DA neurons (Fig. 5A). Apomorphine-induced rotational behavior was observed on days −7 (baseline, prior to 6-OHDA) and on days 7, 14, and 21 after unilateral 6-OHDA injection into the DLS. We found that when compared to Wt female mice, β2-upregulated transgenic female mice demonstrated significantly reduced apomorphine-induced contralateral rotations on days 7, 14, and 21 post-6-OHDA injection (Number of contralateral rotations/15 min: Wt mice day -7, 8.4 ± 1.6; β2-upregulated mice day -7, 11.12 ± 1.61; Wt mice day 7, 178.5 ± 23.1; β2-upregulated mice day 7, 146.94 ± 10.22; Wt mice day 14, 236 ± 31.93; β2-upregulated mice day 14, 163.82 ± 10.97; Wt mice day 21, 233 ± 24.90; β2-upregulated mice day 21, 180.18 ± 11.52; p = 0.024 and F_(1,25)_ = 5.74 with main effect of genotype, based on two-way repeated measures ANOVA) (Fig. 5C). These data show that β2-upregulated female transgenic mice display a significant reduction in 6-OHDA-induced parkinsonian motor deficits when compared to Wt littermates.

**Figure 5.**
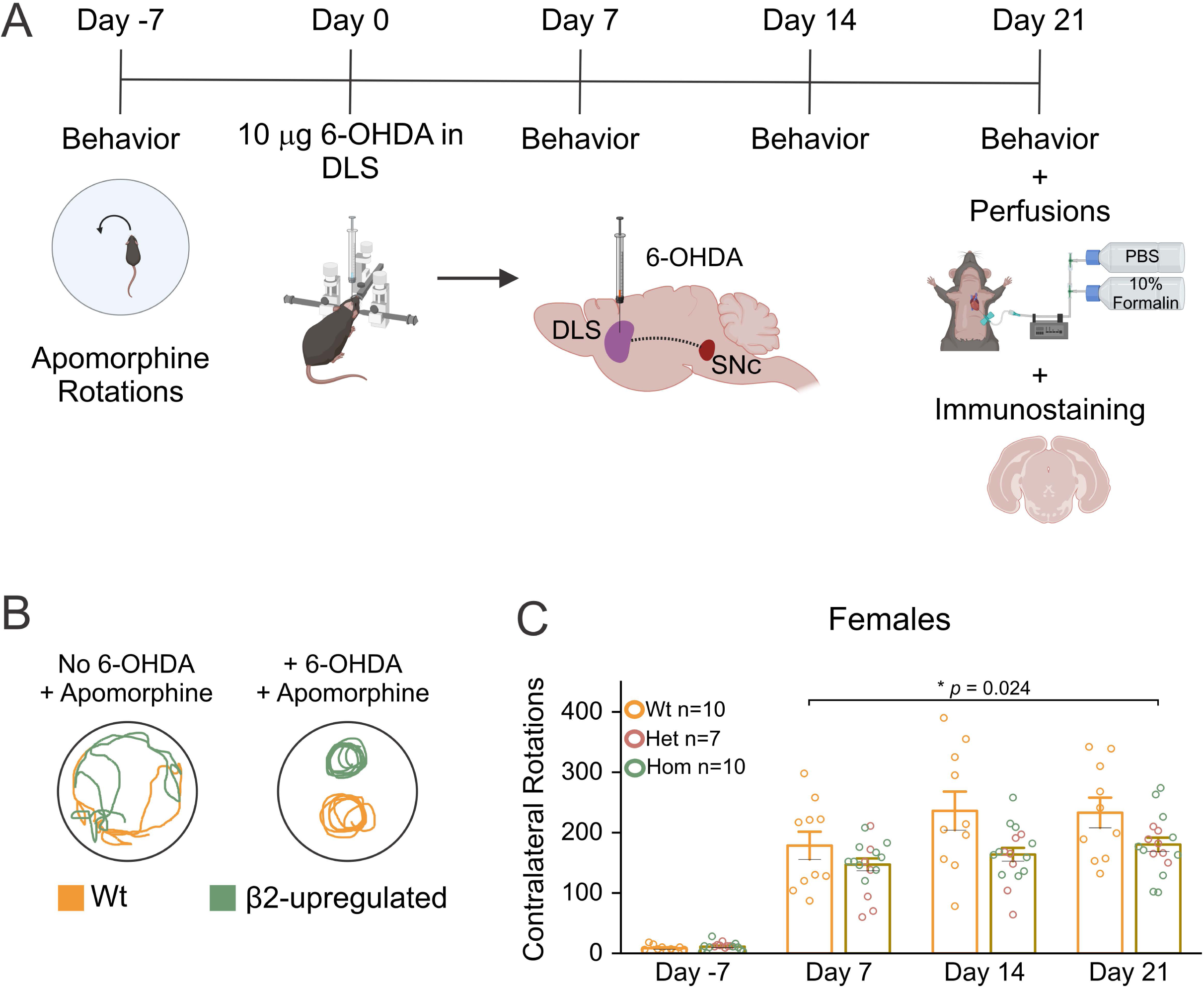
6-OHDA-induced apomorphine rotations are attenuated in β2-upregulated female mice. **(A)** Schematic showing the experimental protocol and timeline for *in vivo* experiments in female mice. Behavioral testing was conducted on days −7, 7, 14, and 21, with unilateral striatal 6-OHDA lesion performed on day 0. Mice were perfused, and brains were extracted for immunostaining on day 21. **(B)** Representative traces of apomorphine-induced contralateral rotations in female Wt and β2-upregulated mice injected with 0.5 mg/kg apomorphine (i.p.) before the 6-OHDA lesion (left) and on day 21 after the lesion (right). **(C)** Bar graphs showing apomorphine-induced rotations on days −7, 7, 14, and 21. n: Wt n = 10 and β2-upregulated n = 17 (Het n = 7, Hom n = 10). p-value is based on a two-way repeated measures ANOVA (genotype*time) for comparisons in number of apomorphine-induced rotations across days after 6-OHDA lesions on days 7, 14, and 21. Error bars represent ± S.E.M. Panel A was created using BioRender.

### Female 6-OHDA lesioned β2-upregulated mice show neuroprotection in the SNc

Having observed a reduction in apomorphine-induced contralateral rotations following unilateral striatal 6-OHDA lesions in β2-upregulated female transgenic mice, we sought to determine if SNc TH+ DA neurons were preserved in female transgenic mice relative to their Wt littermates following exposure to 6-OHDA. At 21 days post-6-OHDA injections, we perfused all the Wt, Het and Hom mice that were assessed for apomorphine-induced rotational behavior (Fig. 5A), and TH immunostaining was performed in 8 evenly spaced midbrain sections obtained from each mouse. To control for differences in TH staining between sections and across mice, the area of SNc TH fluorescence was measured on the lesioned and unlesioned side of each section (Fig. 6A), and a ratio of TH area in the lesioned to unlesioned side was obtained for each section. We found that when compared to Wt mice, the lesioned/unlesioned TH area ratio was ∼34 % greater in β2-upregulated mice (Fig. 6B; Wt: 0.14 ± 0.009, β2-upregulated: 0.21 ± 0.01, *p* = 0.002). To further verify this finding and control for sampling bias in individual midbrain sections versus whole animals, we obtained lesioned/unlesioned side ratios of summed TH area across 8 sections for each mouse. When lesioned/unlesioned TH area ratios were quantified in this way per mouse, β2-upregulated mice showed a ∼2-fold increase in TH area ratios when compared to Wt littermates (Fig. 6C; Wt: 0.13 ± 0.02, β2-upregulated: 0.2 ± 0.03, p = 0.03), thereby confirming that DA neurons in the SNc of β2-upregulated female transgenic mice are more resilient to 6-OHDA-induced neurodegeneration.

**Figure 6.**
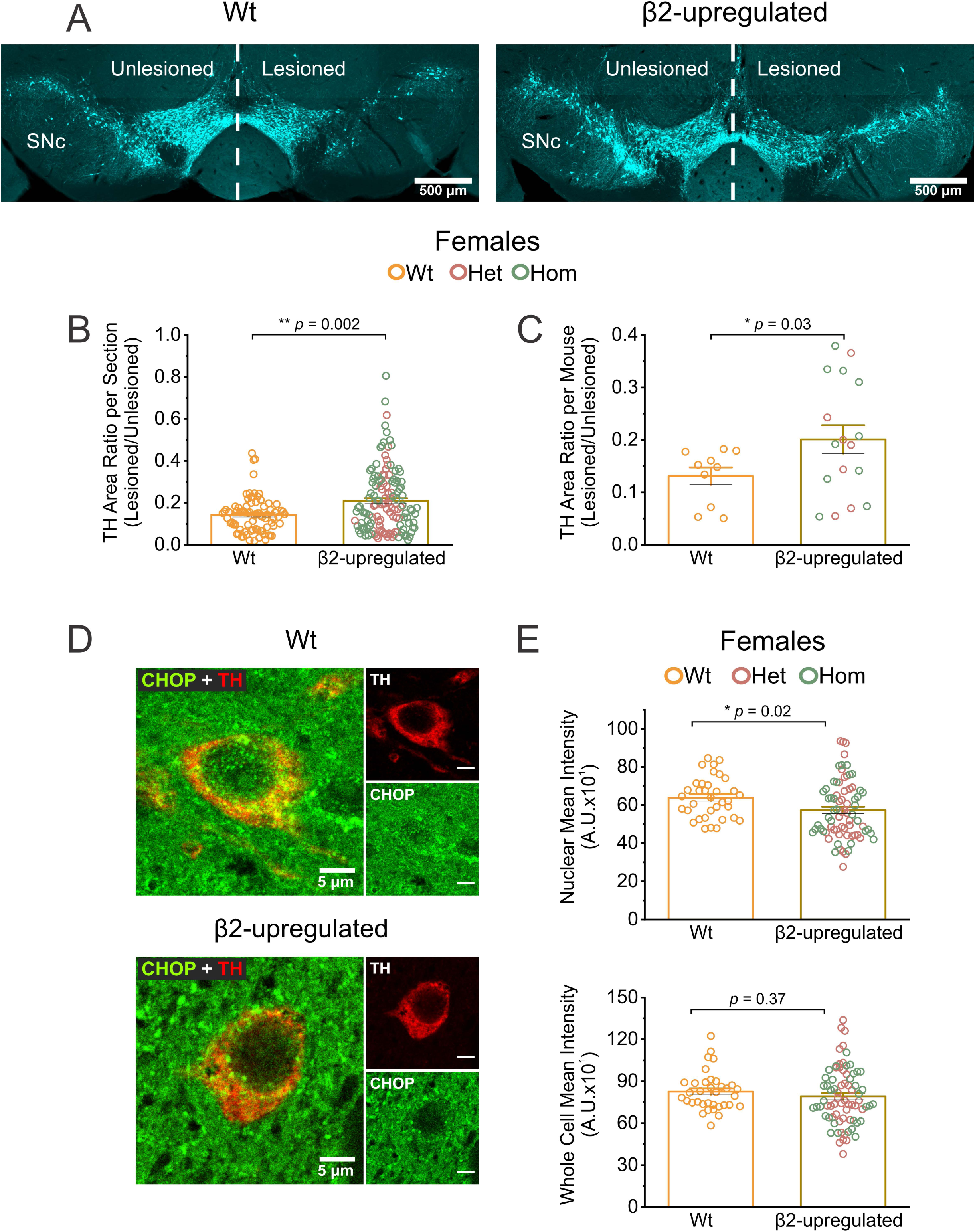
6-OHDA-induced SNc DA neurodegeneration is reduced in β2-upregulated female mice. **(A)** Representative images of midbrain sections from Wt (left) and β2-upregulated (right) female mice. Sections used for analysis showed a clear boundary separating the SNc and VTA. ROIs were manually demarcated for the unlesioned and lesioned midbrain. Scale bar, 500 µm. **(B)** and **(C)** Bar graphs comparing the ratio of TH fluorescence area in lesioned/unlesioned SNc. Data are plotted per section in B or per mouse in C. **(D)** Representative confocal images of DA neurons in lesioned SNc stained for TH (red) and CHOP (green) from Wt (top) and β2-upregulated (bottom) female mice. Scale bar, 5 µm. **e** Bar graphs showing CHOP mean intensity in the nucleus (top) and whole cell (bottom). Sampling for sections B: Wt n = 80; β2-upregulated n = 133 (Het n = 53, Hom n = 80). Sampling for mice in C where 8 midbrain sections were summed per mouse: Wt n = 10; β2-upregulated n = 17 (Het n = 7, Hom n = 10). Sampling for E where each data point represents one cell: Wt n = 36 and β2-upregulated n = 72 (Het n = 39, Hom n = 36). All p-values are based on two-sample t tests and error bars represent ± S.E.M.

### SNc DA neurons in 6-OHDA lesioned β2-upregulated female mice show reduced nuclear CHOP

The finding that SNc DA neurons in female β2-upregulated mice show increased resilience to 6-OHDA mediated cell death (Fig. 6B and 6C) motivated us to ask if when compared to Wt mice, surviving SNc DA neurons in transgenic mice demonstrate reduced activation of CHOP, which is a major mediator of pro-apoptotic ER stress. To do this, we imaged and quantified CHOP fluorescence intensity in surviving SNc DA neurons on the 6-OHDA lesioned side of Wt and transgenic mice (Fig. 6D). When compared to Wt mice, β2-upregulated mice exhibited a significant decrease in nuclear CHOP fluorescence, with no difference across the whole cell between the genotypes (Nuclear CHOP intensity, Wt: 638.86 ± 17.96 and β2-upregulated: 573.61 ± 17.54, *p* = 0.02, two-sample t-test; Whole cell CHOP intensity [AU], Wt: 826.83 ± 23.3 and β2-upregulated: 792.78 ± 23.51, *p* = 0.37, two-sample t-test) (Fig. 6E). Together, these data show that although total CHOP levels in surviving SNc DA neurons were similar between Wt and transgenic mice, CHOP translocation into the nucleus was significantly reduced in β2-upregulated mice. Since CHOP mediates apoptosis via translocation into the nucleus,^26^ these data suggest that β2-upregulated transgenic female mice demonstrate an attenuation of 6-OHDA-mediated apoptotic ER stress in SNc DA neurons.

### Female 6-OHDA lesioned β2-upregulated mice display reduced SNc astrocyte reactivity

Reactive astrocytes are a feature of clinical Parkinson’s disease,^27^ which is thought to further exacerbate disease progression.^28^ In addition, multiple animal models of Parkinson’s disease exhibit elevated expression of astrocytic GFAP in the SNc,^29–31^ suggesting a central role for reactive astrocytes in Parkinson’s disease pathogenesis. We therefore sought to quantify GFAP expression in SNc astrocytes within unlesioned and 6-OHDA lesioned sides of Wt and transgenic female mice at 21 days post-6-OHDA injection (Fig. 7A). Wt mice unilaterally injected with saline were used as a further baseline control to assess changes in SNc astrocyte GFAP expression that could result from stereotaxic surgery. We found significant differences in SNc GFAP intensity between saline-injected and 6-OHDA-injected Wt and β2-upregulated transgenic mice on the unlesioned and lesioned sides. These differences persisted when data were quantified based on either individual sections or averaged between sections to obtain average SNc GFAP intensity per mouse (Fig. 7B and 7C). Specifically, when compared to Wt saline-injected mice, Wt mice with unilateral injection of 6-OHDA into the DLS showed a significant ∼2-fold increase in GFAP intensity in the lesioned as well as the unlesioned SNc (Saline-injected Wt mice: 7.76 x 10^8^ ± 1.95 x 10^7^ A.U per section and 7.76 x 10^8^ ± 3.53 x 10^7^ A.U per mouse on the unlesioned side and 7.64 x 10^8^ ± 2.17 x 10^7^ A.U per section and 7.64 x 10^8^ ± 3.79 x 10^7^ A.U per mouse on the lesioned side; 6-OHDA-injected Wt mice: 1.72x 10^9^ ± 9.57 x 10^7^ A.U per section and 1.72 x 10^9^ ± 1.34 x 10^8^ A.U per mouse on the unlesioned side and 2.1 x 10^9^ ± 1.02 x 10^8^ A.U per section and 2.1 x 10^9^ ± 1.5 x 10^8^ A.U per mouse on the lesioned side; p < 0.0001, ANOVA, followed by post-hoc Tukey). Furthermore, when compared to Wt 6-OHDA-injected mice, β2 6-OHDA-injected mice showed a significant reduction in SNc GFAP intensity in the unlesioned and lesioned sides (6-OHDA-injected Wt mice: 1.72x 10^9^ ± 9.57 x 10^7^ A.U per section and 1.72 x 10^9^ ± 1.34 x 10^8^ A.U per mouse on the unlesioned side and 2.1 x 10^9^ ± 1.02 x 10^8^ A.U per section and 2.1 x 10^9^ ± 1.5 x 10^8^ A.U per mouse on the lesioned side; 6-OHDA-injected β2-upregulated Het and Hom mice: 1.05 x 10^9^ ± 1.0 x 10^8^ A.U per section and 1.09 x 10^9^ ± 1.35 x 10^8^ A.U per mouse on the unlesioned side and 1.47 x 10^9^ ± 1.36 x 10^8^ A.U per section and 1.52 x 10^9^ ± 2.13 x 10^8^ A.U per mouse on the lesioned side; p < 0.0001, ANOVA, followed by post-hoc Tukey). Taken together, these data suggest that β2-upregulated transgenic female mice demonstrate reduced reactivity of astrocytes in the SNc following unilateral 6-OHDA injection into the DLS.

**Figure 7.**
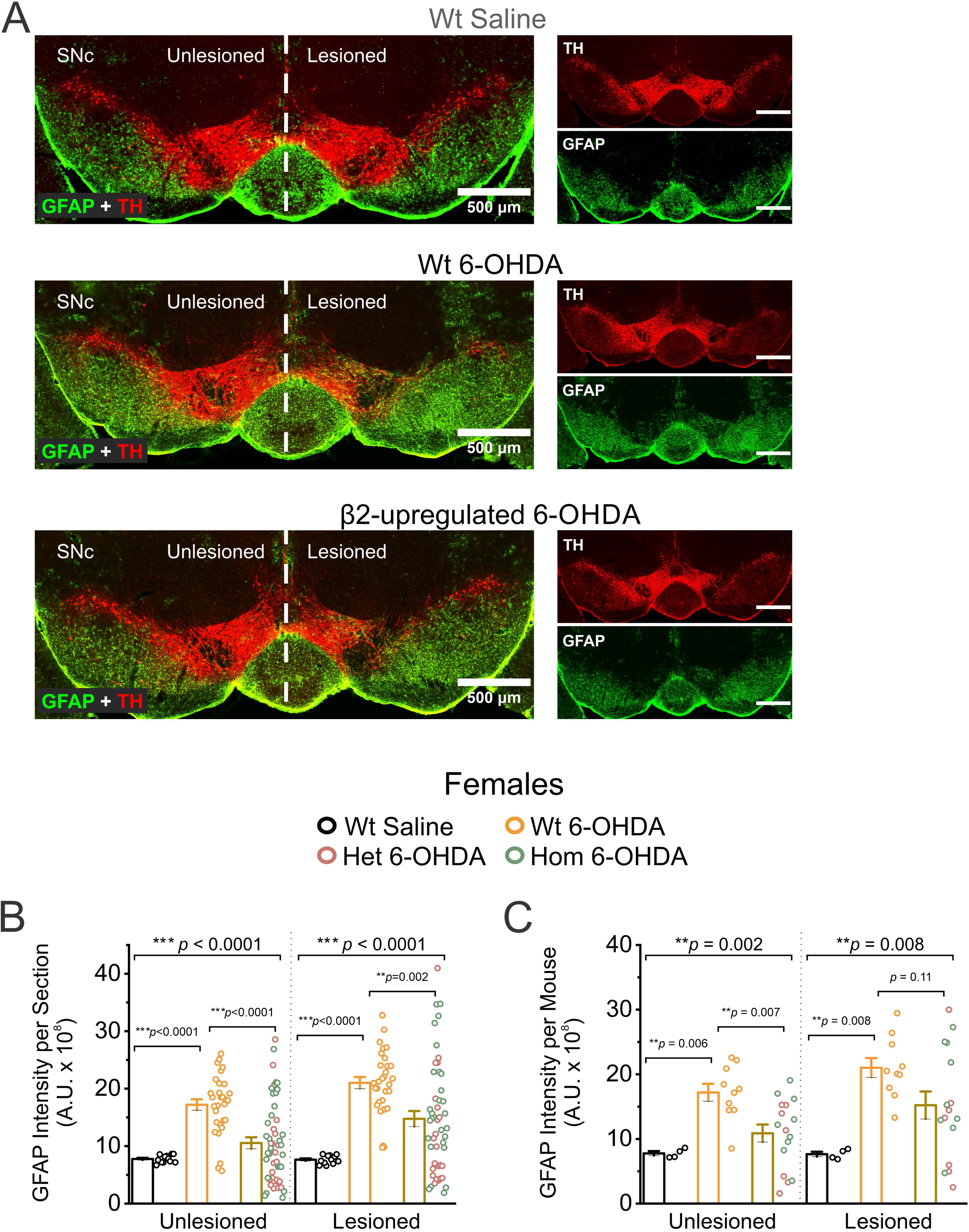
β2-upregulated female mice have reduced astrocytic reactivity in the SNc following 6-OHDA lesion. **(A)** Representative confocal images of midbrain sections stained for TH (red) and GFAP (green) from Wt control saline injected (top), Wt 6-OHDA lesioned (middle), and β2-upregulated 6-OHDA lesioned (bottom) female mice. ROIs were manually demarcated for the unlesioned and lesioned SNc. Scale bar, 500 µm. **(B)** and **(C)** Bar graphs comparing GFAP intensity in the unlesioned and lesioned SNc per section (B) and per mouse (C). Sampling for sections: Wt saline n = 12, Wt 6-OHDA n = 30, β2-upregulated n = 51. Sampling for mice where 3 midbrain sections were averaged per mouse: Wt saline n = 4, Wt 6-OHDA n = 10, β2-upregulated n = 17. All p-values for the unlesioned and lesioned comparisons are derived from a one-way ANOVA comparing each treatment group and error bars represent ± S.E.M.

## Discussion

In this study, we generate and characterize a transgenic mouse line called β2-upregulated mice with genetically-encoded upregulation of β2* nAChRs. We use these novel transgenic mice to ask the important question of whether β2* nAChR upregulation alone, without exposure to nicotinic ligands, reduces SNc DA neuron loss during Parkinson’s disease. β2-upregulated transgenic mice possess the following two mutations within the M3-M4 intracellular loop of β2 nAChR subunits: (**i**) 344-LFL-346 mutated to 344-LFM-346, and (**ii**) 365-RRQR-368 mutated to 365-AAQA-368 (Fig 1). Our choice of these specific mutations is based on a prior study in which we used *in vitro* overexpression of α4 and β2 nAChR subunits in Neuro-2A (N2a) cells to show that these two genetic mutations enhance the exit of α4β2 nAChR pentamers from the ER, thereby mimicking nAChR upregulation without exposure to nicotinic ligands.^11^ Because the mutated 344-LFM-346 motif specifically utilizes Sec24D-containing ERES for exiting the ER,^11,16^ our finding that only female and not male β2-upregulated mice show an increase in the number of Sec24D-ERES (Fig. 2) strongly indicates that only female transgenic mice demonstrate an increase in the genetically-encoded ER exit and upregulation of β2* nAChRs. One potential explanation for the observed sex difference is that female mice could assemble β2* nAChR stoichiometries that are more amenable to ER exit and surface upregulation. This possibility is corroborated by studies demonstrating sex differences in baseline expression of β2* nAChRs in rodents,^32,33^ as well as sex differences in the expression of specific nAChR subunit transcripts during nicotine exposure and withdrawal in rats and humans.^34,35^ Systemically circulating 17-β-estradiol could be another source for the observed sex-specific upregulation of β2* nAChRs in female mice. In this regard, the C-terminus of α4 subunits in mice and humans contain a WLAGMI sequence motif, which is a binding site for 17-β-estradiol^36–38^. Based on this finding, it is possible that the binding of 17-β-estradiol to the α4 C-terminal WLAGMI sequence in female mice not only acts as an allosteric modulator,^38^ but also conformationally stabilizes intracellular α4β2* nAChRs pentamers, thereby causing enhanced exit of β2* nAChRs from the ER in female but not male transgenic mice.

Having found an increase in Sec24D-ERES only in female transgenic mice (Fig. 2), we employed two independent optogenetic assays to assess the extent to which β2* nAChRs in SNc DA neurons are functionally upregulated in female transgenic mice. To do this, we focused on two spatially segregated subcompartments of SNc DA neurons, *viz.*, the somatodendritic compartment and axonal terminals in the DLS. Within the somatodendritic compartment of SNc DA neurons, bath applied ACh caused a significant increase in DhβE sensitive Ca^2+^ influx, measured using GCaMP6f in TH+ dendrites, when compared to Wt littermates (Fig. 3B-D), suggesting functional β2* nAChR upregulation in SNc DA neuron dendrites. Since β2* nAChRs are non-specific cation channels and the upregulated high sensitivity nAChR stoichiometry (α4)_2_(β2)_3_ that is preferentially assembled following the two mutations in the β2 M3-M4 loop^11^ has relatively low permeability to Ca^2+^ ions,^39^ these data suggest that upregulated β2* nAChRs in transgenic mice are functionally coupled to Ca^2+^ channels within SNc DA neuron dendrites.^40^ Indeed, previous studies have shown that β2 nAChR subunits contain sequence motifs that specifically direct nAChR trafficking to dendrites as well as axons,^41^ and that nAChRs can functionally couple to voltage gated calcium channels (VGCCs).^17–20^ Apart from considerations of the mechanistic cause for increases in ACh-evoked dendritic Ca^2+^ in female transgenic mice, the direct demonstration that upregulation of β2* nAChRs in female transgenic mice causes significant increases in Ca^2+^ influx within the dendrites of SNc DA neurons (Fig. 3) could have important implications for understanding the specific role of β2* nAChR upregulation in DA neuron dendrites in the context of chronic nicotine use.

Based on results showing increased ACh-evoked Ca^2+^ influx into the dendritic compartment of SNc DA neurons in female transgenic mice (Fig. 3), we rationalized that upregulated β2* nAChRs at axonal terminals of SNc DA neurons in the DLS would enhance ACh-evoked dopamine release in female transgenic mice.^22–24^ Therefore, as a second assay for functional β2* nAChR upregulation in SNc DA neuron axons, we used live brain slices from female transgenic and Wt littermate mice to quantify bath ACh-evoked dopamine release from SNc DA neuron axonal terminals in the DLS. We found that bath application of ACh causes a small but significant increase in DhβE sensitive striatal dopamine release, as measured using the optogenetic dopamine sensor, GRABDA (Fig. 4D). This result suggests that axonal terminals of SNc DA neurons in the DLS of female transgenic mice do possess upregulated β2* nAChRs. However, based on the relatively small magnitude of DhβE sensitive increase in dopamine release (∼10 to 15%) in transgenic females compared to Wt littermates, we cannot rule out the possibility that upregulation of β2* nAChRs in the somatodentritic compartment of SNc DA neurons (Fig. 3) could alter dopamine synthesis and the releasable pool of dopamine in SNc DA axons. Indeed, experiments using β2 subunit knockout mice have linked the absence of β2* nAChRs to impaired striatal DA release.^23^ In this context, our new transgenic mice could provide an avenue for specifically understanding how somatodentritic nAChR upregulation in DA neurons without the use of nicotinic ligands can affect striatal dopamine release during anxiety-related behaviors.

Having established that female β2-upregulated transgenic mice possess functionally upregulated β2* nAChRs, we used the following four independent measures to assess SNc DA neuron loss in a preclinical model of parkinsonism with unilateral 6-OHDA injection into the DLS (Figs. 5-7): (**i**) Apomorphine-induced contralateral rotations (Fig. 5), (**ii**) Quantification of TH+ SNc DA neurons (Fig. 6A-C), (**iii**) Quantification of nuclear CHOP translocation in SNc DA neurons, which is indicative of proapoptotic ER stress^42^ (Fig. 6D and E), and (**iv**) Quantification of SNc astrocyte reactivity using SNc GFAP expression as a readout (Fig. 7). We injected a high dose of 6-OHDA into the DLS (10 μg), which caused a dramatic increase in apomorphine rotations (Fig. 5C) and ∼85% loss of SNc DA neurons (Fig. 6C). Within the context of this robust preclinical model, female transgenic mice consistently demonstrated a ∼30% reduction in apomorphine rotations (Fig. 5C) and ∼30% preservation of TH+ SNc DA neurons compared to Wt littermates (Fig. 6C). This correlates with the ∼30% upregulation of Sec24D-ERES in SNc DA neurons of female transgenic mice (Fig. 2B), indicating that constitutive β2* nAChR upregulation via Sec24D-ERES does indeed confer resilience to SNc DA neurons against degeneration. As a third readout of neuroprotection, we found reduced translocation of CHOP into the nucleus of SNc DA neurons in female transgenic mice, but we did not observe a general reduction in CHOP levels across the whole cell (Fig. 6D and E). The finding that there was no reduction in CHOP expression across the whole DA neuron raises the possibility that rather than a direct effect of circulating 17-β-estradiol on CHOP promoter activity and CHOP expression levels, as previously reported,^8,43^ the reduction in nuclear CHOP translocation in female transgenic mice (Fig. 6E) likely reflects a downstream effect of reduced ER stress in SNc DA neurons due to increased nAChR exit from the ER of these cells. Future studies will assess the role of circulating 17-β-estradiol on Sec24D-ERES upregulation and neuroprotection in β2-upregulated female transgenic mice. Given the emerging importance of SNc astrocytes in the pathogenesis of Parkinson’s disease,^27–31^ we assessed astrocyte reactivity in the SNc of Wt and transgenic mice as a fourth independent readout of neuroprotection (Fig. 7). Compared to saline-injected mice, Wt mice with unilateral 6-OHDA lesion displayed a robust increase in GFAP expression and astrocyte reactivity in the lesioned as well as the unlesioned SNc, which was significantly attenuated in 6-OHDA lesioned female transgenic mice (Fig. 7B and C). The finding that both sides of the SNc in our unilateral lesion model display astrocyte reactivity is novel and suggests that secreted factors such as cytokines from reactive astrocytes likely cross the midline, resulting in increased astrocytic reactivity on both sides of the midbrain. This finding may be particularly relevant for beginning to understand the role of midbrain astrocyte reactivity in the clinical evolution from unilateral to bilateral symptoms in Parkinson’s disease.

One limitation of this study is that we do not assess behavioral changes or SNc DA neuron loss in male transgenic mice because of the finding that male transgenic mice do not show an upregulation of Sec24D-containing ERES (Fig. 2). In summary, our study demonstrates for the first time that nAChR upregulation alone is sufficient to confer neuroprotection against SNc DA neuron loss. This finding has important implications for developing clinical neuroprotective treatment strategies against Parkinson’s disease via upregulation of endogenous β2* nAChRs in SNc DA neurons without the need for receptor activation. Apart from relevance to Parkinson’s disease treatment, we envision that our newly created transgenic mice will be a valuable resource for understanding the role of β2* nAChR upregulation in addiction, anxiety, depression and cognitive function.

## Data availability

The data that support the findings of this study are available from the corresponding author, upon reasonable request.

## Acknowledgements

We thank the Texas A&M Institute for Genomic Medicine (TIGM) for their help in creating the β2-upregulated transgenic mice using CRISPR-Cas9 technology and for genotyping and maintenance of transgenic mouse colonies. The authors acknowledge the assistance of the Integrated Microscopy and Imaging Laboratory at the Texas A&M College of Medicine (RRID:SCR_021637).

